# Conserved Untranslated Regions of Multipartite Viruses: Natural Markers of Novel Viral Genomic Components and Tags of Viral Evolution

**DOI:** 10.1101/2022.01.16.476546

**Authors:** Song Zhang, Caixia Yang, Jiaxing Wu, Yuanjian Qiu, Zhiyou Xuan, Liu Yang, Ruiling Liao, Xiaofei Liang, Haodong Yu, Fang Ren, Yafeng Dong, Xiaoying Xie, Yanhong Han, Di Wu, Pedro Luis Ramos-González, Juliana Freitas-Astúa, Changyong Zhou, Mengji Cao

## Abstract

Viruses with split genomes are categorized as being either segmented or multipartite according to whether their genomic segments occur in single or different virions. Some complexity will exist, in that inherited “core” vital segments viruses may renew the others once host and environmental alterations keep driving viral evolution. Despite this uncertainty, empirical observations have been made across the split genomes in the untranslated regions (UTRs) on the short or long stretches of conserved or identical sequences. In this study, we describe a methodology that combines RNA and small RNA sequencing, conventional BLASTx, and iterative BLASTn of UTRs to detect viral genomic components even if they encode orphan genes (ORFans). Within the phylum Kitrinoviricota, novel putative multipartite viruses and viral genomic components were annotated using data obtained from our sampling or publicly available sources. The novel viruses, as extensions or intermediate nodes, enriched the information of the evolutionary networks. Furthermore, the diversity of novel genomic components emphasized the evolutionary roles of reassortment and recombination, as well as genetic deletion, strongly supporting the genomic complexity. These data also suggest insufficient knowledge of these genomic components for categorizing some extant viral taxa. The relative conservation of UTRs at the genome level may explain the relationships between monopartite and multipartite viruses and how the multipartite viruses can have a life strategy involving multiple host cells.

**Author summary:** The current workflows for virus identification are largely based on high-throughput sequencing and coupled protein sequence homology-dependent analysis methods and tools. However, for viruses with split genomes, the identification of genomic components whose deduced protein sequences are not homologous to known sequences is inadequate. Furthermore, many plant-infecting multipartite viruses contain conserved UTRs across their genomic components. Based on this, we propose a practical method of UTR-backed iterative BLASTn (UTR-iBLASTn) to explore the components with ORFans and study virus evolution using the UTRs as signals. These shed light on viral “dark matter”—unknown/omitted genomic components of segmented/multipartite viruses from different kingdoms and hosts, and the origins of these components.

## Introduction

Regardless of host range, genome material (DNA or RNA), structure (circular or linear), or polarity (plus, minus, or both), or gene arrangement, a virus genome is constituted by one molecule, so-called monopartite, or more relatively independent parts. These can be segmented if the nucleic acid molecules are assembled into one viral particle or multipartite if these segments are packaged individually into physically separated virions [1]. In monopartite viral genomes, including the circular ones, a linear variable array of genes shows unidimensional orders and positions, but genetic distributions into more spatial arrays make it multidimensional—thus, re-assortments can take place in segmented/multipartite viruses [2]. Whatever the circumstances, viral genomes can acquire autologous or exogenous sequences in a process called “recombination” to generate recombinant progenies among them [3]. Moreover, segmented/multipartite are not rare among families or genera across virus kingdoms [1, 4]. Whether segmented/multipartite viruses are the next evolutionary steps of the monopartite viruses is still indeterminate [5]. Further, their superiorities, either confirmed, i.e., better virion stability [6] or theoretical [4, 5, 7], cannot be neglected. However, genomic segmentation also incurs within- and between-host costs, especially for multipartite viruses [5], despite a multicellular viral lifestyle that permits spatial segregation and possible infection delay of some components in hosts may in part counteract some of the costs [8].

*Kitrinoviricota* is a phylum under the realm *Riboviria*, with its positive-sense RNA viruses that do not infect prokaryotes grouped into a distinct cluster in the polymerase [9]. Of the four classes in this phylum, two are relevant in this study: *Alsuviricetes* and *Flasuviricetes*. The class *Alsuviricetes* includes a single family, the *Flaviviridae*, which was considered to be non-segmented before the finding of Jingmen-related species [10, 11]. Thus far, it seems that plants and fungi are not its natural hosts. Another class comprises three orders, one of which is *Martellivirales*, where viral genome segmentation occurs frequently. This order includes numerous members that infect plants, such as the families *Closteroviridae, Kitaviridae*, and *Virgaviridae*. In addition, a unique family that only infects animals has been specified as the *Togaviridae*.

High-throughput sequencing (HTS) and, homology-dependent annotation (e.g., BLAST search) that underpins HTS, are classical procedures used in viral metagenomic studies for parsing constituents and structures of the virosphere [12–14]. Accordingly, our understanding can flow along with phylogenetic lineages of diverse viruses with their shared genes as mediums [15], but the diversity of “orphan genes” (ORFans) carried by viruses as genetic vehicles remains largely unknown [16]. In addition, ORFans present in unsegmented viruses can be uncovered with sequence recovery of the entire genome, for the “unidimensional.” However, they may escape from notice in segmented/multipartite viruses due to methodology limitations. For example, directly probing the ORFans with BLASTx or similar is unrealistic. As such, a shift in strategy toward BLASTn is feasible when segmented/multipartite viruses share, between their genomic components, conserved sequences of untranslated regions (UTRs), sometimes successive and extensive, that facilitate the analysis [17]. A solution may also come from an inductive analysis of virus-derived small interfering RNAs (vsiRNAs) patterns arising out of virus-host interactions in terms of RNA silencing [18]; in plant hosts sizes of 21- and 22-nt generally predominate among the vsiRNAs [19]. Here, iterative BLASTn, with real-time updated viral UTRs as the database, led to the discovery of many novel multipartite-like viral genomic segments. They have plant viral small RNA characteristics and also show conserved terminal sequences in their own viral entities. With the presence of their equivalents in field samples and homologs in online datasets, their possible importance cannot be ignored.

## Results

### Hands-on operation of iterative BLASTn and its application range

According to our analysis pipelines (**Fig. 1A**), in the first round of processing, the contigs were assembled *de novo* from clean reads (8.38–14.33 Gb) derived from individual RNA sequencing of leaf tissues of plants—ailanthus, apple, camellia, citrus, jasmine, loquat, and paper mulberry, and subsequently subjected to local BLASTx search (e-value cutoff: e-4) against the prebuilt viral non-redundant protein sequences (nr) database (taxonomy ID: 10239). Then, viral-like sequences were collected. The UTRs of the obtained viral sequences and the related viruses from NCBI databases were then constructed into a local database designated as “VUTR” as a target for later BLASTn search (threshold e-value: e-4) of the unexploited contigs; the resulting extra viral UTRs, if any, from the previous step were added into the VUTR for the next step of BLASTn repeatedly until a null outcome was achieved. This analysis was UTR-iBLASTn (UTR-iterative-BLASTn).

**Figure 1.**
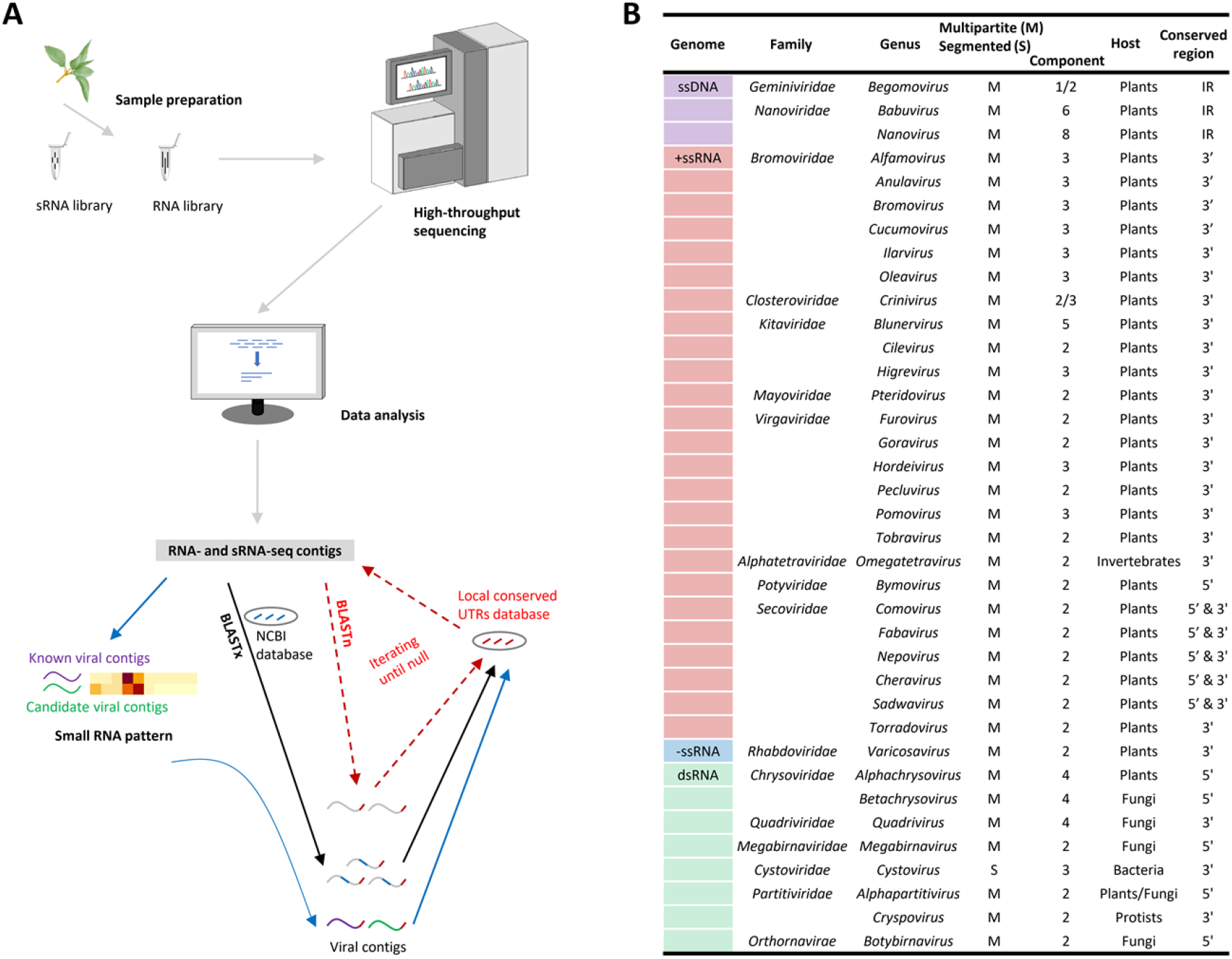
Methodology for the identification of multipartite viral genomic components, based on high-throughput sequencing and viral untranslated regions (UTRs) dependent iterative BLASTn (A), and its most suitable application ranges (B), where the UTRs of a virus are quite conserved (in > 100 nucleotides the sequence identities > 60%). IR: intergenic region. 5’ and 3’: the 5’ and 3’ genomic ends. Consideration of some viruses as multipartite or segmented is based on the phylogenetic information.

Specially selecting the viral UTRs for the analysis opposite to using the entire sequences was considered for potential interferences from non-conserved regions as that may induce additional false-positive results. Furthermore, excluding the data of all other viruses in the following research, 41 viral contigs in total—of which 14 were not recognized by BLASTx due to ORFans (orphan ORFs—open reading frames)—were found to be multipartite-related. They could be divided into eight species in four forms involving two positive-strand (ps) RNA viral orders, three as jivi-like (JVL, 1 species), bluner-like (BNL, 3 species), and crini-like (CNL, 2 species) in the *Martellivirales*, the class *Alsuviricetes*, and one as jingmen-like (JML, 2 species) in the *Amarillovirales*, class *Flasuviricetes*, under the same phylum (Table S1), and a crini-associated satellite virus. Thus, the UTR-iBLASTn as a supplement can extend the relationship complexity of the viral segment (contig) network, upon the BLASTx results, and detect the viral segments in which all of the ORFs are ORFans, named here as orphan segments (ORSs); see Fig. S1. The iteration of BLASTn is obligatory unless, in the assembled data, no contig as a unique pivot is enabling connections with more of the unknowns.

Before 2005, next-generation sequencing (or HTS) was not available for use in virology. Even today, a viral enrichment step is desirable prior to sequencing [20], e.g., double-strand RNA extraction or gradient centrifugation. However, viral nucleic acids and particles of small size and/or low titer could be easily omitted. In this context, the HTS of high sensitivity offers the possibility of including all viral information in single unbiased RNA sequencing with only the removal of ribosomal RNAs, which is the fundamental basis of the UTR-iBLASTn. **Fig. 1B** enumerates viral genera in the most appropriate scope of employing the UTR-iBLASTn, where the taxa are heavily biased toward plant multipartite viruses likely because the great majority of the viruses are hosted in plants.

### Verification of viral orphan segments by molecular cloning, online searching, field investigation, and small RNA typing

With viral-specific primers, the multipartite-like viral sequences were confirmed by RT-PCR amplification, cloning, and Sanger sequencing. Some were completed as full-length genomic RNA by sequencing the terminal sequences. Of the 41 viral contigs, other than partial fragments of the same RNA, all are independent genomic components or satellite RNA; this is likely a result of good sequencing quality and robust sequence assembly. The verified 5’ and 3’ UTR sequences of a virus were regionally conserved whether complete or incomplete.

A search for more orthologs in the NCBI open databases—i.e., the nr (including GenBank), Transcriptome Shotgun Assembly (TSA), Sequence Read Archive (SRA)—using the viral protein and UTR sequences of JML, JVL, and BNL as queries in online or local iBLAST searches found 56 related genomic sequences (44 for TSA, 9 for SRA, and 3 for GenBank) in 12 plant species, semi-unannotated or unannotated before, which potentially belong to 15 viral species (see Table S1 and S2). For the analysis, the SRAs were downloaded and assembled into contigs. A total of 25 of the 56 were ORSs, but the majority were homologs of the ORSs found in the six actual samples (Fig. S1). The minority were four segments related to a novel type of virga-like (VGL) virus from *Rhazya stricta* TSA (RST refers to the mark on the virus) in addition to the aforementioned four types. The remaining 14 species were grouped into JML (three), JVL (nine), and BNL (two). Four field-sampled viral species (three JML and one CNL) have four segments (three and one, correspondently) that are not yet deciphered, apart from the four of RST. We then named the eight ORSs as real-ORSs (R-ORSs) to distinguish them from the other 30 pseudo-ORSs (P-ORSs); R-ORSs would be P-ORSs if their homologs were detected later.

Ignoring any etiological significance that may ascribe symptoms to a virus, we investigated the occurrence of viral non-core RNA molecules (e.g., ORSs) of the eight species (seven are novel) and satellite to determine if the initial genomic combinations persist among samples in the field. Five or more samples per virus were used for statistical purposes, and each sample tested positive by RT-PCR for the replication-associated genomic segment of the virus. A comparison of plant viruses with their animal-infecting counterparts reveals that the generic roles of the additional ORSs in plants are nothing but RNA silencing suppressors used to counter host defenses [21] and movement proteins to direct viral intercellular or systematic trafficking [22]. While they are genomic requisites, none can be found at the protein level for all the JML, JVL, and some BNL viruses. Despite the four peculiar R-ORSs, no ORS or satellite had less than a 20% detection rate in the total tested samples infected by the viruses themselves (Table S3). This suggests the important roles of the broad-sensed ORSs. Furthermore, an RNA2-derived defective RNA (dRNA) of the single infected CNL virus in the sequenced paper mulberry is not an isolated case, since occasional occurrences (6 out of 17) among the other samples were detected. The equal-length sequences of the RNA2 and dRNA showed a 1% nt sequence difference. There are likely to be a large amount of read accumulations of the dRNA in the sequenced sample, because coverages of specific reads on the joined positions of each of four deletions are >3,054 in depth. These suggest the dRNA may be replicable and/or producible in host cells and/or transmissible between hosts.

Besides citrus (1 JVL virus) and paper mulberry (1 CNL virus), other plant samples infected by JML, BNL, and CNL viruses were sequenced to study small viral RNAs (sRNA-seq). Two aspects of the vsiRNAs—the size distribution within 17–27-nt and the 5’ nucleotide preference (5’-nt) among the ACGU—were analyzed to search for a discrepancy between the viruses with the inference that they all infect plants. As anticipated, the results of 21-nt and 22-nt as common peaks of all of the size distributions support that they are plant viruses based on experimental and empirical rules [19]; the genomic segments are clustered mainly by hosts but the variables beyond the host may include virus, dynamic stage, and environment (Fig. S2). The predominant 5’-nt varies randomly with exception of the G, which could be regarded as an evolutionary proclivity of virus-plant RNA silencing interactions. The dynamics, from comparisons between hosts, viruses, or the viral strands (positive or negative) presuppose the targeting specificity of the host RNA-induced silencing complex guided by a complementary small RNA [23]. Thus, it is reasonable to conclude that the viruses have directly interacted with the plant defense system at an intracellular level.

### Metaphylogeny of the Amarillovirales and Martellivirales

The RNA-dependent RNA polymerase (RdRP) gene is shared by viruses in the phylum *Kitrinoviricota*. An RdRP-based tree (Fig. 2) clearly showed, in this phylum, the evolutionary status of the multipartite-like viruses we studied by open-data reanalysis and presented in a demonstration of the UTR-iBLASTn. Moreover, the BNL and CNL viruses were incorporated into clusters representing their own related genera, suggesting close relationships, whereas the branches of the other viruses (JML, JVL, and VGL, respectively) were formed laterally within the Jingmenvirus group (animals), or distally within the families *Togaviridae* (animals) and *Virgaviridae* (plants). These groupings indicated distinct evolutionary paths (**Fig. 2**). We compared these viruses to their closest relatives at the protein level and found that the amino acid sequence identities never exceed 74.2%. Additionally, no significant inconsistencies were observed between the meta-tree and the taxonomically-degraded trees (see below) that were constructed from the RdRP domains with different programs and algorithms.

**Figure 2.**
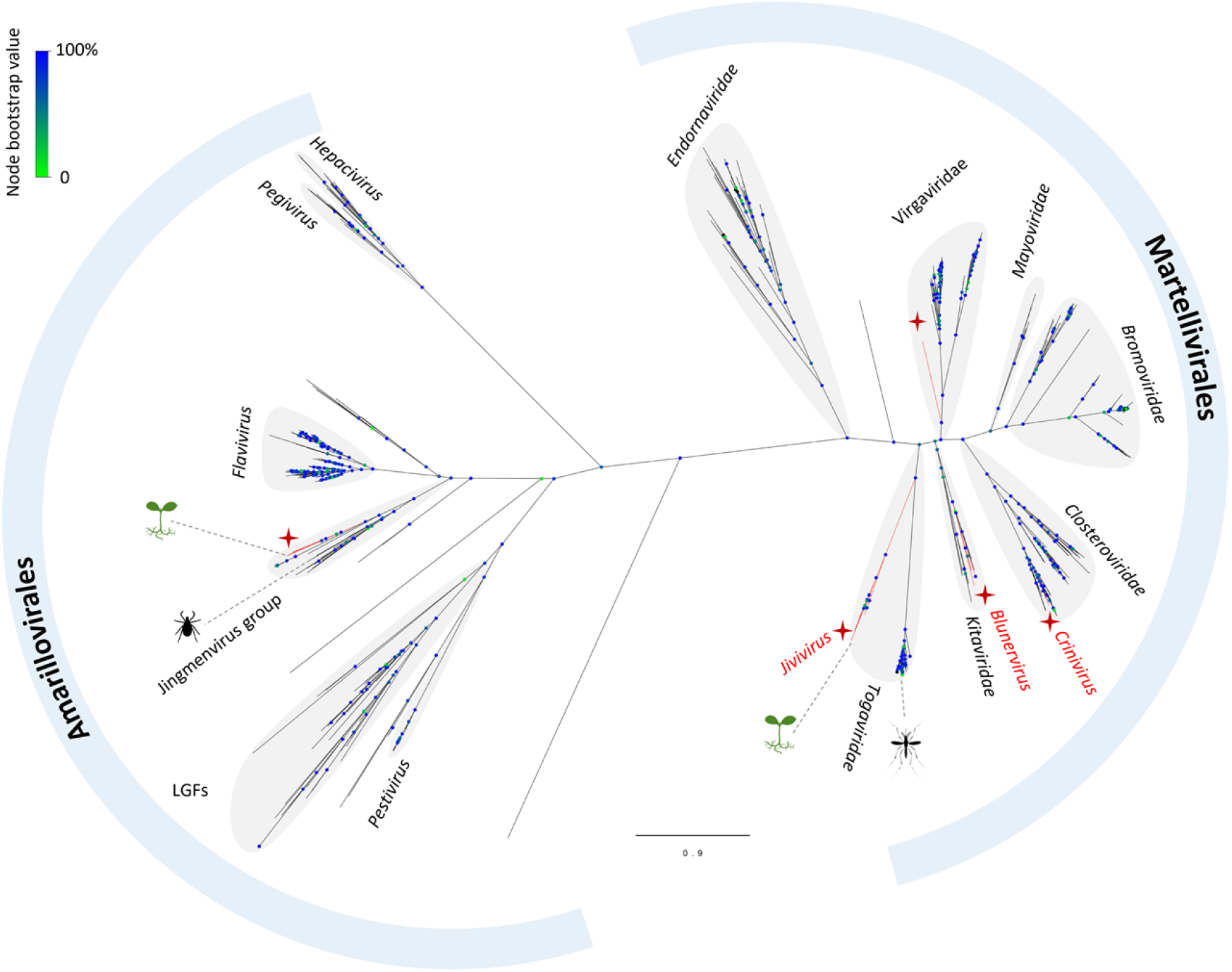
Inferred amino acid sequence alignment of the RdRPs, a phylogram showing evolutionary positions of the new viruses or viral groups (marked in red) under the orders *Amarillovirales* and *Martellivirales*. LGFs: large genome flaviviruses. The splits at the nodes were supported by bootstrap analysis with 1,000 replicates.

### Jingmen- and jivi-like viruses

Considering the host and virus types, the JML and JVL sampled viruses were named ailanthus jingmen-related virus 1 (AJMV1), loquat jingmen-related virus 1 (LJMV1), and citru jivi-related virus 1 (CJVV1). For the two JML viruses from the TSA of *Fagus crenata* (FCT) and *Phalaenopsis equestris* (PET), and one from SRA of grape *Plasmopara viticola* lesions (*Plasmopara viticola* lesion associated Jingman-like virus 1—PVLaJMV1), the genomes comprised two core segments for NS5 (RdRP) and NS3 (DEAD-like helicase superfamily, Hel) proteins plus two P-ORSs for host-specific proteins, with a genomic pattern similar to animal jingmenviruses (**Fig. 3A**). An additional R-ORS in the LJMV1 genomes makes it slightly different, but in the case of AJMV1, one of the P-ORSs was undetectable and two other R-ORSs appeared. This suggested that AJMV1 may represent another genomic pattern. Both the 5’ and 3’ UTRs are conserved whether or not the cases are complete (AJMV1) or partial (LJMV1) genomes (Fig. S3A and B). Independent of the NS5 and NS3 proteins that were analyzed, the phylogenetic trees associated the five plant/fungi JML viruses with those infecting ticks, in a manner of two parallel sub-clusters under the Jingmenvirus group (**Fig. 3B** and Fig. S3C).

**Figure 3.**
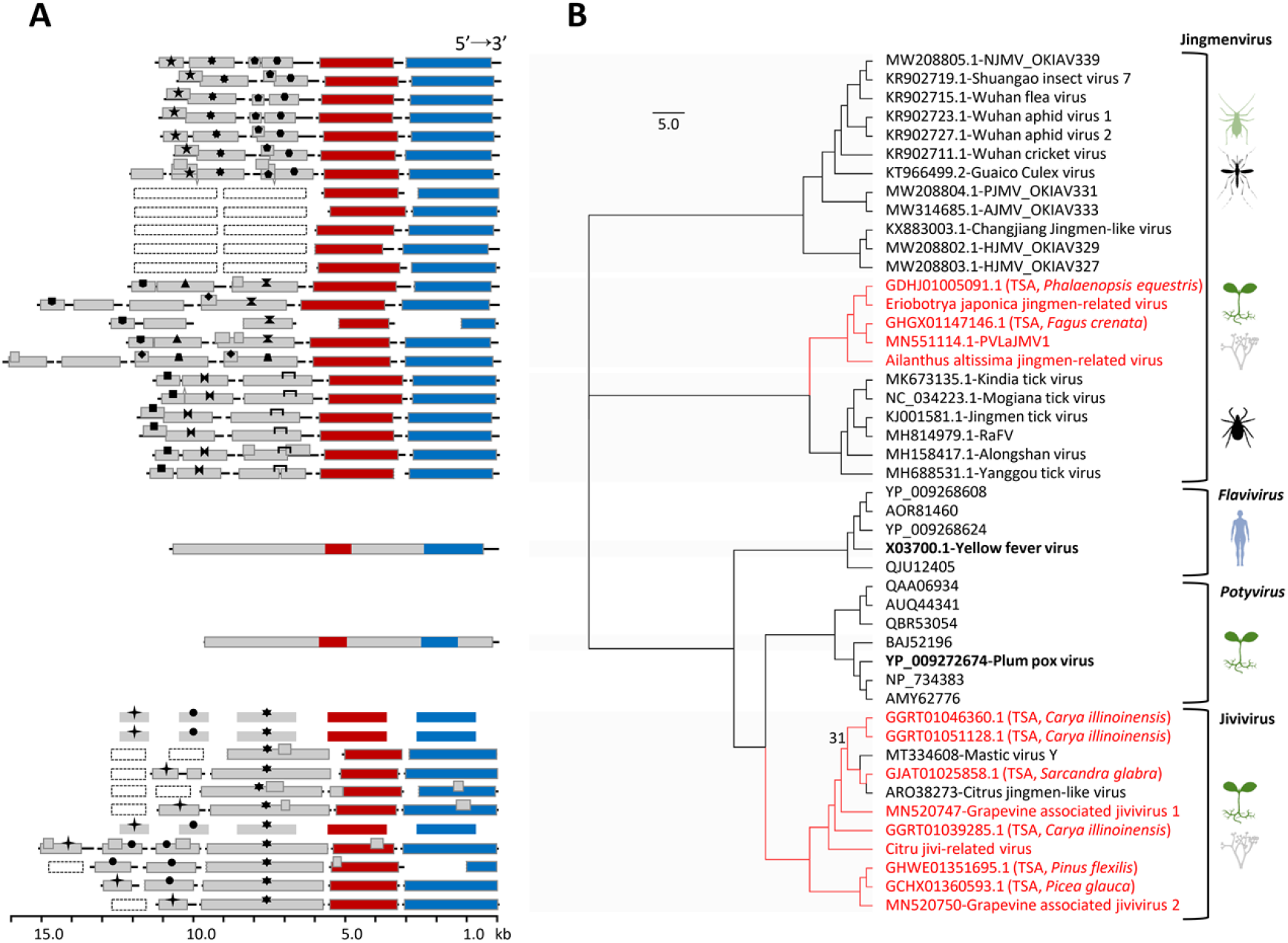
Genomic diagram (*A*) and phylogenetic reconstruction with the LG + G4 model (*B*) of the representative viruses that are homologous in the DEAD-like helicase (Hel) gene according to the results of the BLASTx search using Jiviviruses as queries. The red and blue bars represent respectively the RNA-dependent RNA polymerase (or NS5) and DEAD-like Hel (or NS3); the gray bars with the same symbols indicate protein homologs; the dashed bars are for hypothetical genomic segments. The tree in *B* is for the NS3s. The bootstrap value of a node is not shown unless it had less than 50% value in the test with 1,000 replicates. Red clades represent viruses that have novel genomic components. The viral accession no. is followed by the name or origin, but only one typical virus of the genus *Flavivirus* or *Potyvirus* has been detailed (in bold). These viruses can infect insects (e.g., aphid, mosquitoes), plants and/or fungi, ticks, and mammals (e.g., humans). The virus names have been abbreviated due to size: NJMV, Neuropteran jingmen-related virus; PJMV, Psocopteran jingmen-related virus; AJMV, Arachnidan jingmen-related virus; HJMV, Hemipteran jingmen-related virus; PVLaJMV1, Plasmopara viticola lesion associated Jingman-like virus 1; RaFV, Rhipicephalus associated flavi-like virus.

The only JVL virus sampled was CJVV1, and 10 relatives were found on NCBI. These were citrus jingmen-like virus (or citrus virga-like virus, published) and mastic virus Y (MaVY, semi-annotated): three in TSA of *Carya illinoinensis* (CITs) as coinfections; two in the SRAs of grape *Plasmopara viticola* lesions (grapevine associated jivivirus 1 and 2, GaJV1 and GaJV2); three in the TSAs of *Pinus flexilis* (PFT), *Picea glauca* (PGT), and *Sarcandra glabra* (SGT). CJVV1 (hexapartite). Amongst them, four JVL viruses possess five typical genomic segments for methyltransferase with the helicase 1 superfamily (Mtr-Hel), RdRP, DEAD-like Hel, and two P-ORSs, while the rest (≤ pentapartite) are seemingly genomically incomplete (**Fig. 3A**). Furthermore, the conserved sequences in the UTRs of CJVV1 are sharply reduced in comparison to the JML viruses (Fig. S4A). With variable of protein domains (Mtr, Hel, and RdRP) that used for analysis does not affect the local topology of the trees formed, where the Jivivirus group of plant/fungus origins was always located in the same branch as the mosquito-borne family *Togaviridae* that exclusively infests animals (Fig. S4B–D). However, in the DEAD-like Hel tree (**Fig. 3B**), this group is most closely related to the monopartite families *Potyviridae* (the phylum *Pisuviricota*, plants) and *Flaviviridae* (the phylum *Kitrinoviricota*, animals).

### Bluner-, crini-, and virga-like viruses

Four or fewer RNA segments are associated in genomes of the families *Kitaviridae, Closteroviridae*, and *Virgaviridae*. With the discovery of the new ORSs, we increased the upper boundary to five for all. Typical of the genus *Blunervirus* are four genomic RNAs encoding for Mtr-Hel, Hel-RdRP, the SP24 with three unknown proteins, and MP. From camellia, we previously isolated a variant of the tea plant necrotic ring blotch virus (TPNRBV) named TPNRBV-Ca1; a new P-ORS was also found in SRA of the tea (*Camellia sinesis*). The unusual genomes lack the recognizable MP, which is replaced by new types of homologs (P-ORFs) that were identified in the two BNL viruses named ailanthus blunervirus 1 (AiBV1) and apple blunervirus 1 (ApBV1), and the TSA of *Paulownia tomentosa* (PTT). In addition, both viruses contained a supernumerary RNA—of Mtr-Hel-RdRP for the former and Hel for the other. The phylograms in **Fig 4A, C**, and **D** show *Blunervirus* evolution in the genes of Mtr, Hel, and SP24 (or CP), where the three viruses in a clade are separated from the others, which is consistent with their distinction in the genome aspect (**Fig. 4B** and Fig. S6A–C).

**Figure 4.**
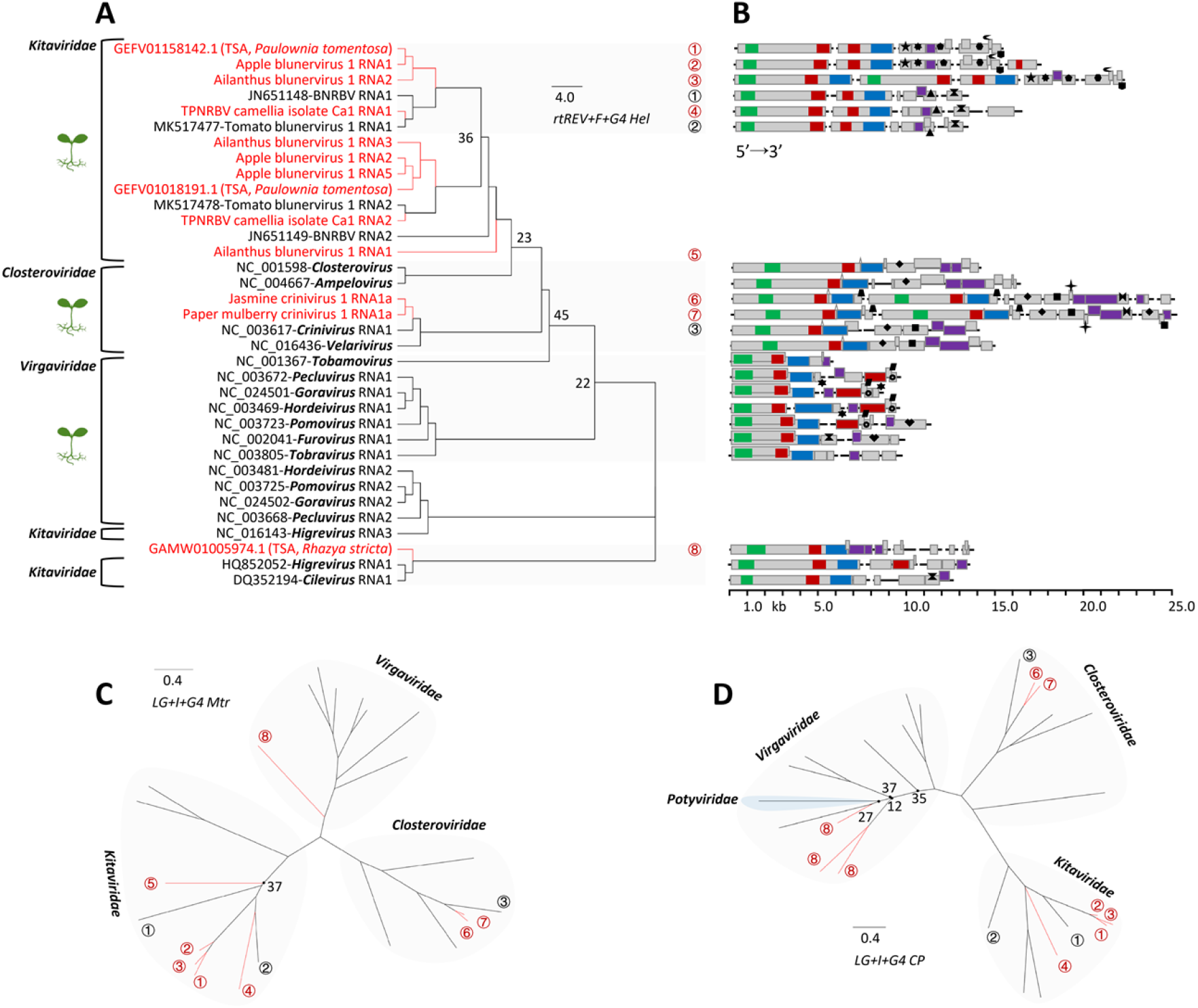
Phylogenetic analysis of the helicase (Hel) protein sequences of the plant-infecting viruses and genera/family that have been confirmed to be closely related in a tree (data not shown) for the order *Martellivirales (A)* and the corresponding viral genomic originations (*B*). The methyltransferase (Mtr) and coat protein (CP) genes were also used for viral phylogenies, as in *C* and *D*, respectively. The tree models adopted are shown in italics, and the novel viruses are indicated in red. With bootstrapping in 1,000 replicates, the node support value is shown if less than 50%. A circled number represents a virus or its RNA. For the genomes, Mtr, RNA-dependent RNA polymerase (RdRP), helicase (Hel), and CP are indicated by green, red, blue, and purple bars, respectively, and not defined regions are represented by gray bars. The protein homologs are signed identically. Abbreviated viral name: BNRBV, Blueberry necrotic ring blotch virus.

Two CNL viruses that were obtained separately from two plant species were named jasmine crinivirus 1 (JCV1) and paper mulberry crinivirus 1 (PMCV1). Like other members in the genus *Crinivirus*, JCV1 and PMCV1 have RNA1 for Mtr-RdRP, and slightly smaller RNA2 for two different heat shock proteins (HSPs), coat protein (CP), and a mirror (CPm), without taking into account other minor genomic ORFs (**Fig. 4B**). This is markedly different in that JCV1 and PMCV1 both have two homologous RNA1s, and JCV1 possesses an additional R-ORS while PMCV1 has a surviving dRNA as a deletion of the RNA2 (Fig. S6A and B). JCV1 was also associated by a satellite virus RNA (abbr. name JCVaSV), which is not singular for criniviruses but unique in a possibly pentapartite genomic concept (Fig. S6C). Accordingly, conservation signals were mainly detected in the 3’ UTRs of both viruses. In the phylogenetics analysis (**Fig. 4A, C**, and **D**), evidence of Mtr, Hel, and CP genes all indicate that the two viruses belong to the genus *Crinivirus*.

The family *Virgaviridae* appears in two or three genomic RNA segments with Mtr-Hel, RdRP, and CP (sole and always alone); but the VGL virus from TSA of *Rhazya stricta* (RST) is a different situation where Mtr-Hel-RdRP and three CPs were integrated into a single RNA, in association with other four small potential genomic RNAs, all of which are R-ORFSs (**Fig. 4B**). The RNAs of the RST are highly identical in the sequences of the 3’ UTRs (Fig. S7). The Mtr, Hel, and RdRP seem phylogenetically connected between the families *Kitaviridae* and *Virgaviridae* (**Fig. 2** and **Fig. 4A** and **C**), and the CPs appear subordinate to the latter (**Fig. 4D**).

### Gene repeats, horizontal gene transfer (HGT), and UTR-phylogeny and -networking

In the forms of independent components within a genome (nine cases) and ORFs in a segment (three cases), gene duplications of less than 97.7% amino acid sequence identities were frequently observed for the 23 viral species in Mtr, Hel, RdRP, CP, and ORSs (Table S2). The extreme cases of a single gene replicated three times were found in AiBV1 and ApBV1, both of which were verified by genomics and in RST (if not chimeric). Aside from the ApBV1-Hels, of which one lacked some cardinal motifs (Fig. S8), the AiBV1-Hel and RST-CP gene repeats appeared to be complete in basic functions. At times, only one of the duplicated P-ORSs (i.e., RNA5s of CJVV1 or PFT, and PMCV1-RNA1s) had a unique non-viral protein domain (CDD e-value < e-10) while the other protein regions were homologous between them. This likely occurred because only some of the original genomic RNA had undergone the HGT [24] by recombination or insertion, but both kinds were retained (Table S2). The AiBV1 included in the *Blunervirus* genus had an RNA1 (Met-Hel-RdRP) related to the possibly bipartite and tripartite genera in the same family, which may suggest another mechanism whereby the duplications are generated—genetic deletion (**Fig. 4A–D**). We propose an evolutionary model with ApBV1 as an example to show historical time nodes at which the Met-Hel-, Hel-RdRP-, and Hel-encoding RNA originated from the partial RNA of Met-Hel-RdRP (Fig. S9). To examine this possibility, we reconstructed the evolutionary tracks of the family *Kitaviridae* in the Hel and its 3’ UTR concomitant, both of which may have derived from the same duplication event (**Fig. 4A** and Fig. S10A and B). Finally, phylogenetic congruence of the Hels and 3’ UTRs is consistent with the result of different genetic deletions of the ancient common genomic RNA. The networking of the UTRs (Fig. S10C) based on local reciprocal BLASTn reinforced this view because, in AiBV1, RNA1 and other RNAs are related, and in ApBV1, the RNAs (1, 2, 5) are correlated with Hels.

## Discussion

To anchor the viral genomic components occupied by ORFans in data flows, our efforts were based upon several earlier concepts [17, 25]. This work has resulted in the UTR-iBLASTn method—which examines the data of many putative multipartite viruses representing two classes in the phylum *Kitrinoviricota*—from which several novel components were discovered. Its application can potentially be expanded to encompass more groups of viruses from various hosts (Fig. 1B), even monopartite species that may be similar to the VGL virus whose RNA2-RNA5 are R-OFSs. We obtained insight into this method: the crucial point is that HTS and assembly generated viral contigs with UTRs being as full as possible to include core informative sites. Regarding this, the strategies below may be helpful to obtain satisfied HTS contigs: 1) to choose symptomatic individuals for sampling, 2) to deploy the appropriate sequencing method (e.g., long reads of 150 nt or longer, and rRNA-depleted RNA-seq for a broad spectrum of viral genomes and RNA derivatives), and 3) to enlarge data size per sequencing.

The existence of flavi-related multipartite-like viruses possibly from plant/fungus hosts have been recognized previously but inadequate information impeded labeling and taxonomic clarity [26–28]. The samples studied here and retrieved from the NCBI databases (GenBank, TSA, or SRA), show that some JML viruses exhibit unambiguous plant virus attributes in vsiRNAs. However, neither plants nor fungi could be eliminated for the jivi-related viruses, as there is no conflict between commonality and graft-transmissible in plants and parasites in plant-fungal endophytes [29]. Furthermore, genomics and phylogenetics analyses indicated that the plant JML viruses descended from a single origin with viruses infecting animals, but the jivi-grouped viruses had multiple origins that likely arose from reassortment or recombination (RNA1s–3s) and the HGT (fungal and bacterial xenologs: RNA5s) may derive from distantly related taxa (*Flaviviridae*, *Potyviridae*, and *Togaviridae*). In a similar case when restricting the scope of phylogeny to the order *Martellivirales*, some newly discovered or annotated viruses may also serve as evolutionary bridges between the families *Kitaviridae* and *Virgaviridae* or genera of the former. Thus, to comprehend viral diversity, both the abundance and mobility of the existing genetic pools and the ability to recombine new genes from other organisms must be considered.

It remains unclear if segmented/multipartite or monopartite forms originated first, but evidence favors the former [4, 30, 31]. There is yet a lack of phylogenetic support on this issue and both forms possess unique advantages and costs [5]. In light of this, transitions between the two forms can occur [6], because natural selection does not rule out either from within a range of viruses that share a common ancestry. This implies parallel evolution. The situation of AiBV1, however, indicates that it is more plausible that monopartite genomes—at least for this given *Kitaviridae* lineages—are the predecessors from which multipartite ones arose. If multipartite/segmented originated first, then it would be difficult to explain why Hel of AiBV1-RNA1 has not phylogenetically skewed to either from the RNA2 and 3. Moreover, the segmentation of the original RNA by selective partial deletions could explain the three Hel paralogs from different but congenetic ApBV1 RNAs (Fig. S9). As previously demonstrated [30]—and with observations on the long-lasting persistence of PMCV1-dRNA (Fig. S11)—deletions of genomic RNA that bring about defective RNA to be complementary as a functional complete systems could be a driving force for viruses to create and update their segmented/multipartite genetic architectures.

The plant multipartite virus—faba bean necrotic stunt virus (FBNSV, genus *Nanovirus*)—is a complement of viral genomic components from different host cells where they accumulate, likely through the trafficking of functional materials like mRNA and protein products that simulate host operation [8]. Some of these products that are only effective on viruses themselves (e.g., RdRP for replication) may be exported from original hijacked cells, and delivered by host transportation to cells adjacent or even distant, to search/identity inactive cognate viral genomic components and act upon them. While this scenario is reasonable, one checkpoint is how these adventitious viral products can be precisely targeted even if we exclude the complex process of guiding them to specific endocellular compartments or organelles containing the objectives [32] as well as assume their high intracellular abundance contrary to the genome formulas [33]. In addition, monopartite and segmented/multipartite viral genomes arrange their untranslated regions principally into both ends (liner genome) or a large intergenic region (circular type) pertaining to a concentration of regulatory elements or signals for stability, transcription, replication, and packaging [34]. For a segmented virus, the packaging of virions can be executed through a programed process initiated from a certain single component with its UTRs (also essential for replication) and medicated by between-components networking—a simple yet subtle design [35]. Multipartite viruses may abandon the direct component interactions when the situation is complex as a spatial scale elongated beyond single cells, with the cost of indispensably equipping each component with all of the necessary instruments. The extensive conserved sequences in the UTRs (Fig. 1B) can be a manifestation of such a cost as multipurpose recognition sites in support of viral multicellular network reconstruction and circulation. Accordingly, indirect evidence comes from some plant satellites that preserve analogous UTRs to their helper viruses, e.g., JCVSV and JCV1 (Fig. S6C), likely to remain compatible.

The components in a multipartite genome appear to be complementary and organized. This rule can also cover the subviral plant satellites as viral parasites, from which their helper viruses may gain pathogenicity. The relationship may become interdependent [36], despite the satellite viruses encoding CP for themselves and being partially independent. Similarly, endogenous or exogenous nucleic acid molecules and their duplications may also embed in a viral genome in the course of coevolution. This may be how some viral ORSs originated. However, triple repetitions of viral genes are present beyond the fully functional system. There may be collateral subsystems under the genome with each working with one copy, or simply they may coordinate in, or reinforce, the same function.

In summary, using the UTR-iBLASTn method and our or open-sourced data, we identified new multipartite viruses or viral genomic components that fill evolutionary gaps or blanks and emerge as new phylogenetic twigs. These data deepened our understanding of viral genome segmentation, expanded our knowledge of virus diversity and their split genomes, and broadened our view of the genomic components of variation among species and higher taxa.

## Materials and Methods

### Sample collection, HTS, BLASTx, iBLASTn, and networking

Leaf samples with virus-like symptoms were collected from seven plant species during field studies in three provinces of China: Chongqing (camellia, citrus, loquat, and paper mulberry), Liaoning (ailanthus and apple), and Yunnan (jasmine). The total RNA of each sample (from one plant) was extracted using the EASY spin Plus Complex Plant RNA Kit (Aidlab, Beijing, China). The RNA purity, concentration, and integrity were evaluated using a Nanodrop (Thermo Fisher Scientific, Cleveland, OH, USA), Qubit 3.0 (Invitrogen, Waltham, MA, USA), and Agilent2100 (plant RNA Nano Chip, Agilent, Santa Clara, CA, USA), respectively. The ribosome RNA was depleted by the RiboZero Magnetic Kit (Epicenter, Madison, WI, USA), and a library was then built using a TruSeq RNA Sample Prep Kit (Illumina, San Diego, CA, USA). Furthermore, RNA-seq was conducted by Beijing Genomics Institution (BGI) using the BGI500 platform set 100 bp for the length of paired-end (PE) reads, by Mega Genomics (MG, Beijing, China) using an Illumina HiSeq X-Ten platform (PE 150 bp), or by Berry Genomics Corporation (BGC, Beijing, China) with an Illumina NovaSeq 6000 platform (PE 150 bp). The generated HTS data were assembled using the CLC Genomics Workbench 11 (Qiagen, Hilden, Germany) after the removal of adaptors and low-quality reads, as well as the host reads, with the draft genomes of related plant species used as references if available. Subsequently, local BLASTx and iBLASTn were performed using the fast-speed DIAMOND [37] and BLAST-2.12.0+, respectively. For online data, the verified viral sequences were subjected to NCBI-BLASTn and -BLASTx searches against the TSA and nr/nt databases, and new sequences were used as new queries until there were no additional results. The SRA database was also tested with BALSTn, and the TSA and SRA databases were restricted to higher plants (taxid:3193). Only results with an e-value < e-4 were used for further analysis. The NCBI searches were accessed in 2019 and 2020. The selected sequences were loaded into a final test using local iBLASTn-2.12.0+. Finally, the ORSs were determined and the P-OFSs and R-ORSs differentiated. All of the resulting data were networked by the Cytoscape 3.8.0 [38] with Atribute Circle Layout that arranged these by the RNA names and removing nonsignificant edges (BLAST e-value > e-4) between nodes.

### Recovery and confirmation of viral sequences

Specific primer pairs were designed at appropriate positions of the viral contigs to amplify overlapping fragments using the CLC Genomics Workbench 11 and the Primer Premier 5.0 (Premier Biosoft, Palo Alto, CA, USA). Thereafter, a one-step reverse transcription-PCR (RT-PCR) assay was conducted with a PrimeScript kit (Takara, Tokyo, Japan), and Viral genomic terminal sequences were determined using commercial 5’ and 3’ RACE (rapid amplification of cDNA ends) kits (Invitrogen, Waltham, MA, USA). The PCR products were gel-purified using the Gel Extraction Kit (OMEGA Bio-Tec Inc., Doraville, GA, USA) and cloned on pEASY-T1 Vectors (TransGen, Beijing, China) using competent cells. Five clones of each amplicon were fully sequenced with primer in both directions (Tsingke, Chengdu, China), and the output sequences were *de novo* assembled in the SeqMan program (DNAStar, Madison, WI, USA).

### Field investigation

Virus-specific detection primers were designed in the conserved coding regions using the same programs mentioned above. To avoid high viral heterogeneity between different samples, we collected plants within a limited area in an orchard, garden, or field, less than 0.01 km^2^ in size. The replication of ORSs relied on the RdRP: therefore, we first detected the associated RNAs in samples by the one-step RT-PCR, and the positive samples were further tested for the presence of ORSs.

### Viral small RNA profiles

In sRNA-seq, total sRNA was extracted from leaf tissues with the EASYspin Plant microRNA Extract kit (Aidlab, Beijing, China). The sRNA library was constructed using a TruSeq Small RNA Sample Prep Kit (Illumina, San Diego, CA, USA). Sequencing platforms were selected among Illumina Hiseq2500 platform (MG, Beijing, China), NextSeq CN500 (BGC, Beijing, China), and BGI500 (BGI, Beijing, China). The resulting sRNA was processed to remove useless sequences, and the remaining reads directly mapped on the HTS contig sequences as references using the CLC Genomics Workbench 11. Through the statistical analysis, putative viral contigs with sRNAs distribution characteristics in the size and 5’-nt similar to the known viral sequences were collected. The data of the distributions of vsiRNAs from all of the multipartite-like viral contigs were normalized and visualized with the pheatmap package 1.0.12 in R [39].

### Sequence analysis

The open reading frames (ORFs) of viral nucleotide sequences and conserved domain in protein sequences inferred from ORFs were found using the NCBI ORF finder (https://www.ncbi.nlm.nih.gov/orffinder) and CDD (https://www.ncbi.nlm.nih.gov/Structure/cdd/wrpsb.cgi) servers, respectively. The ORSs were reanalyzed using the DIAMOND-BLASTx with the local nr database. These steps were accomplished to identify known genes/segments and the HGTs from non-viral organisms. Within-group relationships of the ORSs were reestablished by local BLASTx-2.12.0+ with the threshold e-value e-4 for each JML, JVL, CNL, and BNL-VGL viral group to differentiate between the P-OFSs and R-ORSs. The viral amino acid sequences were compared, and identities were calculated using the CLC Genomics Workbench 11.

### Phylogenetic analysis

For the meta-tree of the phylum *Kitrinoviricota*, representative viral RdRP sequences of the order *Amarillovirales* were selected as previously accomplished and retrieved from NCBI by the Batch Entrez server (https://www.ncbi.nlm.nih.gov/sites/batchentrez); viral genomes of the order *Martellivirales* (taxonomy ID: 2732544) were obtained from the NCBI databases; the RdRP sequences were identified using the Batch CD-Search server (https://www.ncbi.nlm.nih.gov/Structure/bwrpsb/bwrpsb.cgi) and extracted by the Strawberry Perl (5.32.1.1) using a script. All of the RdRPs were aligned by MAFFT v7.471 [40], using the E-INS-i algorithm, and then trimmed by trimAl 1.2rev57 [41], with the automated1 algorithm to remove poorly aligned regions. Then, the phylogenetic relationships of the remaining alignments were computed by the FastTree 2.1.11 [42] with the LG + CAT model. The other viral protein sequences were analyzed by the same methods, but the results were processed by the IQ-TREE 1.6.12 [43], with the best-suited model automatically selected by the component tool according to the Bayesian Information Criterion (BIC) with 1,000 bootstrap replicates. All the tree files were determined by the FigTree v1.4.4 (http://tree.bio.ed.ac.uk/software/figtree/).

### Evolution of the UTRs

For the family *Kitaviridae* and the RST, the 3’ UTRs of each virus were separately aligned instead of directly by the MAFFT (E-INS-i) and trimmed by the trimAl (automated1). This was done to obtain conserved signals specific to each virus and to avoid false phylogenetic connections between any two viruses. All of the alignments of the conserved sequences were merged and analyzed through the workflows of the same MAFFT algorithm, the IQ-TREE with the recommended model, and 1,000 bootstraps; FigTree was used to draw and modify the phylogram. Local BLASTn-2.12.0+ with the parameter not filtering low-complexity regions was executed within the viral UTRs that are used as both queries and targets. The yFiles Radial Layout in the Cytoscape was used to display networks of the UTRs as nodes with the e-values as edges (e-4 as the cutoff for visible).

## Data Availability

All viral sequences obtained from this study are available in NCBI databases with accession numbers OL344024–OL344048 or in the Data S1. The sequencing datasets generated in this work can be provided for reasonable requests. All other study data are included in the main text and/or supporting information.

## Acknowledgments

This research was supported by the National Key R&D Program of China (2019YFD1001800), National Natural Science Foundation of China (32072389), 111 Project (B18044), Earmarked Fund for China Agriculture Research System (CARS-26-05B) and Chongqing Postgraduate Research and Innovation Project (CYB21133). We thank LetPub (www.letpub.com) for its linguistic assistance during the preparation of this manuscript.

## Author Contributions

SZ and MC designed the experiments; SZ, YC, JW, YQ, ZX, LY, RL, XL, HY, FR, XX, YH, YD, and MC provided materials, conducted experiments, and analyzed the data; SZ, PLRG, FAJ, JF, CZ, and MC discussed the results and prepared the manuscript.

## References

1. Sicard A, Michalakis Y, Gutiérrez S, Blanc S. The strange lifestyle of multipartite viruses. PLoS pathogens. 2016;12(11):e1005819.

2. Varsani A, Lefeuvre P, Roumagnac P, Martin D. Notes on recombination and reassortment in multipartite/segmented viruses. Current opinion in virology. 2018;33:156–66.

3. Simon-Loriere E, Holmes EC. Why do RNA viruses recombine? Nature Reviews Microbiology. 2011;9(8):617–26.

4. Lucía-Sanz A, Manrubia S. Multipartite viruses: adaptive trick or evolutionary treat? NPJ Systems Biology and Applications. 2017;3(1):1–11.

5. Michalakis Y, Blanc S. The curious strategy of multipartite viruses. Annual Review of Virology. 2020;7:203–18.

6. Ojosnegros S, Garcia-Arriaza J, Escarmis C, Manrubia SC, Perales C, Arias A, et al. Viral genome segmentation can result from a trade-off between genetic content and particle stability. PLoS genetics. 2011;7(3):e1001344.

7. Lucía-Sanz A, Aguirre J, Manrubia S. Theoretical approaches to disclosing the emergence and adaptive advantages of multipartite viruses. Current opinion in virology. 2018;33:89–95.

8. Sicard A, Pirolles E, Gallet R, Vernerey M-S, Yvon M, Urbino C, et al. A multicellular way of life for a multipartite virus. Elife. 2019;8:e43599.

9. Koonin EV, Dolja VV, Krupovic M, Varsani A, Kuhn JH. Create a megataxonomic framework, filling all principal taxonomic ranks, for realm Riboviria. 2019.

10. Qin X-C, Shi M, Tian J-H, Lin X-D, Gao D-Y, He J-R, et al. A tick-borne segmented RNA virus contains genome segments derived from unsegmented viral ancestors. Proceedings of the National Academy of Sciences. 2014;111(18):6744–9.

11. Ladner JT, Wiley MR, Beitzel B, Auguste AJ, Dupuis II AP, Lindquist ME, et al. A multicomponent animal virus isolated from mosquitoes. Cell host & microbe. 2016;20(3):357–67.

12. Shi M, Lin X-D, Chen X, Tian J-H, Chen L-J, Li K, et al. The evolutionary history of vertebrate RNA viruses. Nature. 2018;556(7700):197–202.

13. Shi M, Lin X-D, Tian J-H, Chen L-J, Chen X, Li C-X, et al. Redefining the invertebrate RNA virosphere. Nature. 2016;540(7634):539–43.

14. Roossinck MJ, Martin DP, Roumagnac P. Plant virus metagenomics: advances in virus discovery. Phytopathology. 2015;105(6):716–27.

15. Koonin EV, Dolja VV, Krupovic M, Varsani A, Wolf YI, Yutin N, et al. Global organization and proposed megataxonomy of the virus world. Microbiology and Molecular Biology Reviews. 2020;84(2):e00061–19.

16. Yin Y, Fischer D. Identification and investigation of ORFans in the viral world. BMC genomics. 2008;9(1):1–10.

17. Li C-X, Shi M, Tian J-H, Lin X-D, Kang Y-J, Chen L-J, et al. Unprecedented genomic diversity of RNA viruses in arthropods reveals the ancestry of negative-sense RNA viruses. elife. 2015;4:e05378.

18. Aguiar ERGR, Olmo RP, Paro S, Ferreira FV, de Faria IJdS, Todjro YMH, et al. Sequence-independent characterization of viruses based on the pattern of viral small RNAs produced by the host. Nucleic acids research. 2015;43(13):6191–206.

19. Hull R. Plant virology: Academic press; 2013.

20. Wu Q, Ding S-W, Zhang Y, Zhu S. Identification of viruses and viroids by next-generation sequencing and homology-dependent and homology-independent algorithms. Annual review of phytopathology. 2015;53:425–44.

21. Burgyán J, Havelda Z. Viral suppressors of RNA silencing. Trends in plant science. 2011;16(5):265–72.

22. Navarro JA, Sanchez-Navarro JA, Pallas V. Key checkpoints in the movement of plant viruses through the host. Advances in virus research. 2019;104:1–64.

23. Mi S, Cai T, Hu Y, Chen Y, Hodges E, Ni F, et al. Sorting of small RNAs into Arabidopsis argonaute complexes is directed by the 5’ terminal nucleotide. Cell. 2008;133(1):116–27.

24. Gilbert C, Cordaux R. Viruses as vectors of horizontal transfer of genetic material in eukaryotes. Current opinion in virology. 2017;25:16–22.

25. De La Peña M, García-Robles I. Intronic hammerhead ribozymes are ultraconserved in the human genome. EMBO reports. 2010;11(9):711–6.

26. Chiapello M, Rodríguez-Romero J, Ayllón M, Turina M. Analysis of the virome associated to grapevine downy mildew lesions reveals new mycovirus lineages. Virus evolution. 2020;6(2):veaa058.

27. Chiapello M, Rodríguez-Romero J, Nerva L, Forgia M, Chitarra W, Ayllón MA, et al. Putative new plant viruses associated with Plasmopara viticola-infected grapevine samples. Annals of Applied Biology. 2020;176(2):180–91.

28. Matsumura EE, Coletta-Filho HD, Nouri S, Falk BW, Nerva L, Oliveira TS, et al. Deep sequencing analysis of RNAs from citrus plants grown in a citrus sudden death-affected area reveals diverse known and putative novel viruses. Viruses. 2017;9(4):92.

29. Silva JMF, Fajardo TVM, Al Rwahnih M, Nagata T. First report of grapevine associated jivivirus 1 infecting grapevines in Brazil. Plant Disease. 2021;105(2):514–.

30. García-Arriaza J, Manrubia SC, Toja M, Domingo E, Escarmís C. Evolutionary transition toward defective RNAs that are infectious by complementation. Journal of virology. 2004;78(21):11678–85.

31. Ramos-González PL, Santos GFd, Chabi-Jesus C, Harakava R, Kitajima EW, Freitas-Astúa J. Passion fruit green spot virus genome harbors a new orphan orf and highlights the flexibility of the 5’-end of the RNA2 segment across cileviruses. Frontiers in microbiology. 2020;11:206.

32. Heinlein M. Plant virus replication and movement. Virology. 2015;479:657–71.

33. Sicard A, Yvon M, Timchenko T, Gronenborn B, Michalakis Y, Gutierrez S, et al. Gene copy number is differentially regulated in a multipartite virus. Nature communications. 2013;4(1):1–8.

34. Dreher TW. Functions of the 3’-untranslated regions of positive strand RNA viral genomes. Annual review of phytopathology. 1999;37(1):151–74.

35. Sung P-Y, Roy P. Sequential packaging of RNA genomic segments during the assembly of Bluetongue virus. Nucleic acids research. 2014;42(22):13824–38.

36. Zhou X. Advances in understanding begomovirus satellites. Annual review of phytopathology. 2013;51:357–81.

37. Buchfink B, Xie C, Huson DH. Fast and sensitive protein alignment using DIAMOND. Nature methods. 2015;12(1):59–60.

38. Smoot ME, Ono K, Ruscheinski J, Wang P-L, Ideker T. Cytoscape 2.8: new features for data integration and network visualization. Bioinformatics. 2011;27(3):431–2.

39. Kolde R, Kolde MR. Package ‘pheatmap’. R package. 2015;1(7):790.

40. Katoh K, Standley DM. MAFFT multiple sequence alignment software version 7: improvements in performance and usability. Molecular biology and evolution. 2013;30(4):772–80.

41. Capella-Gutiérrez S, Silla-Martínez JM, Gabaldón T. trimAl: a tool for automated alignment trimming in large-scale phylogenetic analyses. Bioinformatics. 2009;25(15):1972–3.

42. Price MN, Dehal PS, Arkin AP. FastTree 2–approximately maximum-likelihood trees for large alignments. PloS one. 2010;5(3):e9490.

43. Nguyen L-T, Schmidt HA, Von Haeseler A, Minh BQ. IQ-TREE: a fast and effective stochastic algorithm for estimating maximum-likelihood phylogenies. Molecular biology and evolution. 2015;32(1):268–74.

